# Gut microbiota transplantation drives the adoptive transfer of colonic genotype-phenotype characteristics between mice lacking catestatin and their wild type counterparts

**DOI:** 10.1101/2022.02.24.481402

**Authors:** Pamela González-Dávila, Markus Schwalbe, Arpit Danewalia, René Wardenaar, Boushra Dalile, Kristin Verbeke, Sushil K Mahata, Sahar El Aidy

**Author notes:** Shared first author (Pamela González-Dávila, Markus Schwalbe). Shared correspondence: Sushil K Mahata; Sahar El Aidy.

## Abstract

The gut microbiota is in continuous interaction with the intestinal mucosa via metabolic, neuro- immunological, and neuroendocrine pathways. Disruption in levels of antimicrobial peptides produced by the enteroendocrine cells, such as catestatin, has been associated with changes in the gut microbiota and imbalance in intestinal homeostasis. However, whether the changes in the gut microbiota have a causational role in intestinal dyshomeostasis has remained elusive. To this end, we performed reciprocal fecal microbial transplantation in wild-type mice and mice with a knockout in the catestatin coding region of the chromogranin-A gene (CST-KO mice). Combined microbiota phylogenetic profiling, RNA sequencing, and transmission electron microscopy were employed. Fecal microbiota transplantation from mice deficient in catestatin (CST-KO) to microbiota-depleted wild-type mice induced transcriptional and physiological features characteristic of a distorted colon in the recipient animals, including impairment in tight junctions, as well as an increased collagen area fraction indicating colonic fibrosis. In contrast, fecal microbiota transplantation from wild-type mice to microbiota-depleted CST-KO mice reduced collagen fibrotic area, restored disrupted tight junction morphology, and altered fatty acid metabolism in recipient CST-KO mice. This study provides a comprehensive overview of the murine metabolic- and immune-related cellular pathways and processes that are co-mediated by the fecal microbiota transplantation and supports a prominent role for the gut microbiota in the colonic distortion associated with the lack of catestatin in mice. Overall, the data show that the gut microbiota may play a causal role in the development of features of intestinal inflammation and metabolic disorders, known to be associated with altered levels of catestatin and may, thus, provide a tractable target in the treatment and prevention of these disorders.

## Introduction

It is well-established that the gut microbiota has an essential role in the development and maintenance of the human physiology by sustaining homeostatic processes such as gut barrier function (Shi et al., 2017), host immunity (Jiao et al., 2020), energy metabolism (A. M. Martin et al., 2019), and neuropsychological behaviours (El Aidy et al., 2016). Disruptions in the intimate interactions between the gut microbiota and the host are positively correlated with pathologies such as inflammatory gastrointestinal diseases, metabolic diseases, and neuropsychiatry disorders (Araújo et al., 2017; El Aidy et al., 2014; Heijtz et al., 2011) The pro-hormone chromogranin-A (CgA) is proteolytically processed to several biologically active peptides including Catestatin (CST: hCgA352-372) (Lee et al., 2003; S K Mahata et al., 1997). CST consists of 21 amino acids and acts as an inhibitor of the catecholamine secretion through activation of nicotinic cholinergic receptors in cultured cells and mice adrenal glands (S K Mahata et al., 1997; Sushil K. Mahata et al., 2000, 2003). Several metabolic and inflammatory disorders have been linked to altered levels of CST (Chen et al., 2019; Kojima et al., 2018; Muntjewerff et al., 2022; Ying et al., 2018b). For example, the administration of human CST in mouse models of chronic inflammation, such as colitis ameliorated the intestinal pro-inflammatory parameters including macrophage function, reduction of pro-inflammatory cytokines and pathways such as interleukin 6 (IL-6), interleukin 1β, tumour necrosis factor α (TNF-α) and signal transducer and activator of transcription 3 (STAT3) dependent pathway (Eissa et al., 2018; Rabbi et al., 2014, 2017). Moreover, the phenotype of mice with selective deletion of the CST-coding region of the *ChgA* gene (CST-KO mice) was shown to display obesity, insulin resistance, hypertension, macrophage infiltration, hyperadrenergic state, as well as high levels of pro-inflammatory cytokines (Ying et al., 2018a, 2021), and more recently an IBD-like phenotype, including intestinal permeability (Muntjewerff, Tang, et al., 2021).

Recently, we and others have shown that CST-KO mice display altered gut microbiota composition compared to their wild-type counterparts (González-Dávila et al., 2021; Muntjewerff, Lutter, et al., 2021). In particular, CST treatment reduced the abundance of *Staphylococcus* and *Turicibacter* in CST-KO and WT mice, while *Alistipes*, *Akkermansia*, and *Roseburia* were significantly increased in the CST-KO group (González-Dávila et al., 2021).

Additionally, levels of the short chain fatty acids (SCFAs), butyrate and acetate, were significantly increased in CST-KO mice treated with CST (González-Dávila et al., 2021). Analogously, supplementation of CST-KO mice with CST restored paracellular intestinal epithelial permeability, reversed inflammation, and fibrosis, all of which are characteristic of CST-KO mice (Muntjewerff, Tang, et al., 2021). This led us to hypothesize that the altered gut microbiota may play a key role in developing the disrupted CST-KO-associated phenotypes. One of the most frequently used experimental approaches to study the causal role of the gut microbiota in gut dysbiosis-related diseases is the fecal microbiota transplantation (FMT) (Bokoliya et al., 2021). Thus, in this study, we performed transplantation of the perturbed microbiota from CST-KO mice to microbiota-depleted WT mice and vice versa in an attempt to study the contribution and the mechanisms by which the gut microbiota may contribute to the phenotype observed in the CST-KO mice.

## Results

### Reciprocal fecal microbiota transfer between CST-KO and wild-type mice harbouring distinct microbial populations

Recently, we and others have shown that mice with a *C*ST knock-out (CST-KO) have a significantly altered gut microbiota composition compared to their wild-type (WT) counterparts (González-Dávila et al., 2021). Additionally, previous studies reported significant differences in gastrointestinal morphology, mucosal immune function, and gut permeability in CST-KO mice (Muntjewerff, Tang, et al., 2021). Since CST-KO mice displayed altered gut microbiota composition, we hypothesized that the microbiota may play a substantial role in causing these differences. To unravel this causality, we performed reciprocal fecal microbiota transplantation (FMT), where C57BL/6 WT mice (n=12) were orally gavaged with a fecal microbiota suspension of CST-KO mice and vice versa (**Figure 1A; methods section; Supplementary Figure 1A**). The transplanted microbiota was allowed to recolonize the gut of the recipient mice for 14 consecutive days. Fecal pellets collected from CST-KO^FMT-WT^ and WT^FMT-CST-KO^ recipient mice and their controls (n=12, per group) were used for amplicon sequencing of the V3-V4 regions of the bacterial 16S gene.

**Figure 1:**
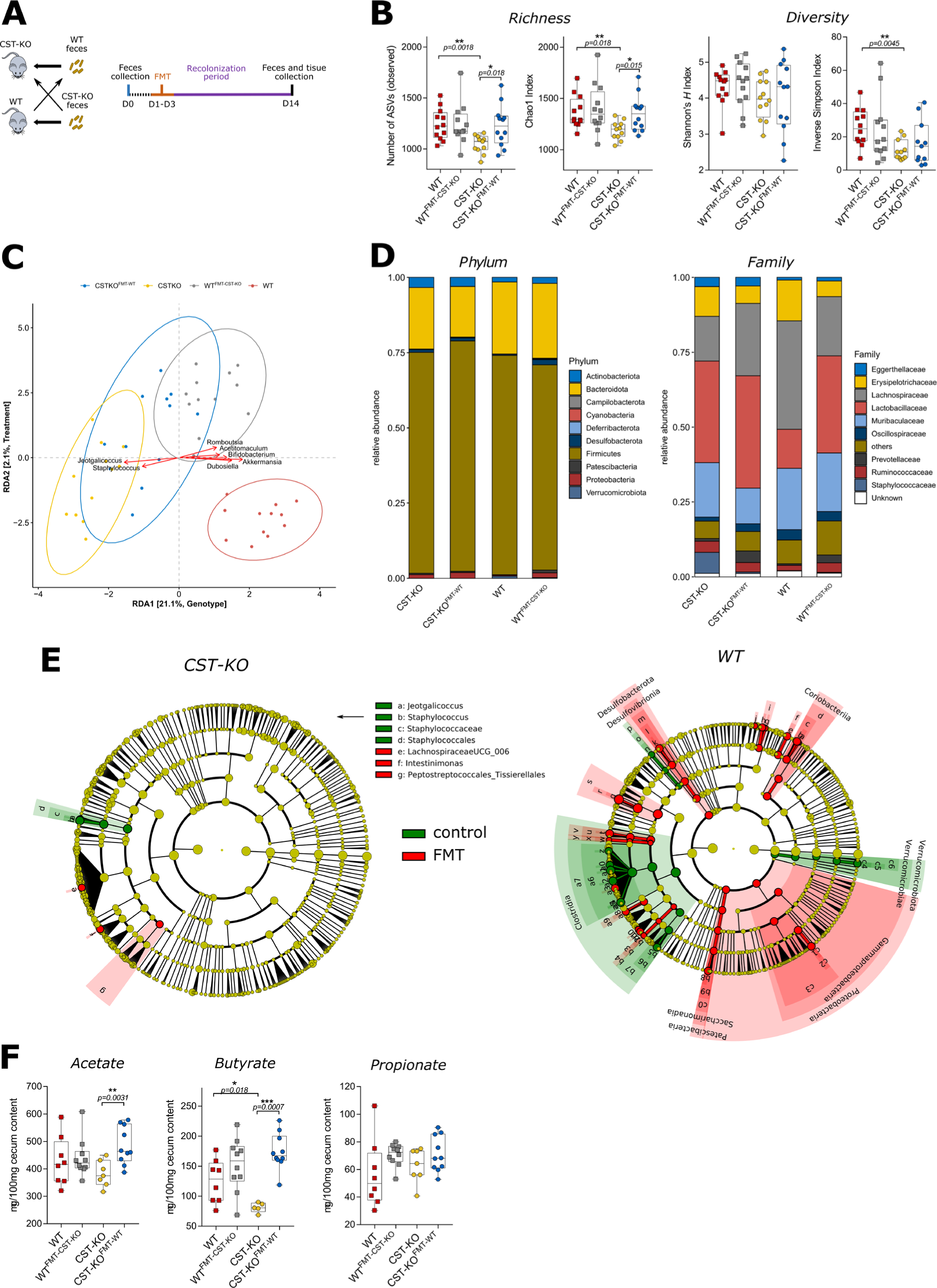
Altered microbiota composition following cross-FMT in CST-KO and WT mice. (**A**) Experimental strategy with timeline. (**B**) Alpha diversity was assessed using different metrics. In CST-KO mice, FMT increased the bacterial richness (observed ASVs and Chao1 index) and diversity (inverted Simpson’s index). Significance was tested with an unpaired *Mann-Whitney* test. Boxes represent the median with interquartile range, and whiskers represent the maxima and minima. (**C**) Genotype and FMT treatment constrained redundancy analysis (RDA) on genus collapsed abundances. Arrows indicate the association of taxa with samples, with the length being a proxy for the strength of the association. Significant separation of clusters and contribution of the variables to the variance of the RDA was tested with Permutational Multivariate ANOVA (PERMANOVA) and revealed significant (p<0.001) effects of FMT, in both CST-KO and WT mice; yellow – CST-KO, blue – CST-KO FMT, grey – WT FMT and red – WT. (**D**) Relative abundance of the present phyla and families (**E**) Cladograms of a LEfSe analysis (Linear discriminant analysis Effect Size) comparing microbiota changes upon FMT in CST-KO and WT mice. The cladogram indicates the microbiota composition represented by rings with phyla in the outermost ring and genera in the innermost ring. The green color represents taxonomic levels enriched in control animals, while the red color represents enrichment in the FMT groups. Each circle is a member within that level. (**F**) Concentrations of cecal acetate, butyrate and propionate of untreated mice (CST-KO, n=7; WT, n=8) and mice after FMT (CST-KO FMT, n=10; WT FMT, n=10). Data were analyzed using a 2-tailed paired *t*-test.

As a general exploratory analysis, principal component analysis (PCA) was performed on genus level collapsed data and showed significantly distinct clustering of the CST-KO and WT before and after FMT. Particularly, the CST-KO and WT donors clustered separately, while the recipients, CST- KO^FMT-WT^ and WT^FMT-CST-KO^ mice, were overlapping into a middle point, indicating that the transplant induced changes in the microbiota composition in both genotypes **(Supplementary Figure 1B)**. Next, microbial richness was assessed by the observed number of amplicon sequence variants (ASVs) and Chao1 index. Interestingly, the richness levels were restored in CST-KO^FMT-WT^, but not in WT^FMT-CST-KO^ mice **(Figure 1B)**. Contrary to the richness scores, the diversity indexes determined by Shannon’s *H* and inverted Simpson’s index, showed no significant changes between the recipient groups **(Figure 1B)**. Analogous to our previous findings (González-Dávila et al., 2021), the CST-KO mice consistently showed decreased richness and diversity compared to their WT counterparts. Taken together, the data infers that FMT induced changes in the microbiota composition in both genotypes, with a significant effect in restoring the richness of the microbiota community in CST-KO mice to the WT level.

To correct for any residual variation in the data, genotype and treatment-constrained (FMT) redundancy analysis (RDA) was performed. In agreement with the PCA **(Supplementary Figure 1)**, distinct clustering was visible and both constraints also had a significant influence on the model (p<0.001, determined by ANOVA-like permutation test), explaining 21.1% (genotype) and 2.1% (FMT treatment) of the variation (**Figure 1C**). RDA determined which bacterial taxa were associated with each group of mice. Focusing on the strongest associations, the largest differences were determined by the genotype. In agreement with our previous findings (González-Dávila et al., 2021), among others, *Akkermansia, Dubosiella, and Bifidobacterium* showed the strongest association with WT along RDA1 while *Jeotgalicoccus and Staphylococcus* were associated with CST-KO mice.

To further identify which bacterial taxa were transferred by the FMT, pairwise comparisons of bacterial abundances were performed between the donor and recipient groups for each genotype. Focusing on the phylum level, *Bacteroidota* decreased, while *Firmicutes* increased in relative abundance, although not significant, in CST-KO^FMT-WT^ mice, similar to the microbiota of WT mice (**Figure 1D**). In contrast, in WT ^FMT-CST-KO^ mice, the opposite effects were observed, where *Firmicutes* and *Verrucomicrobiota* significantly decreased in relative abundance, while *Bacteroidota*, *Actinobacteriota*, and *Patescibacteria* were significantly increased, again similar to the microbiota of CST-KO mice **(Supplement Excel Sheet 1)**. On the family level, the most significant changes were observed for the increased abundance of *Peptostreptococcaceae*, *Staphylococcaceae*, *Defluviitaleaceaea*, as well as UCG-010 in CST-KO^FMT-WT^ mice, all of which decreased in abundance in WT^FMT-CST-KO^ mice **(Supplement Excel Sheet 1)**.

To complement our analysis and to search for predictors at all taxonomic levels including genera, LEfSe was employed (Linear discriminant analysis Effect Size, Segata et al. 2011) **(Figure 1E).** Consistent with the RDA, the main discriminant feature separating the groups in CST-KO mice were species from the genera *Jeotgalicoccus* and *Staphylococcus*, while CST-KO^FMT-WT^ mice were represented with *Lachnospiraceae UCG_006*, *Intestinimonas*, and *Peptostreptococcaceae*. Notably, compared to the CST-KO groups, alterations in the microbiota composition were more pronounced in the WT mice, where these animals were enriched in species from the class *Clostridia* and *Verrucomicrobiota*, while the WT^FMT-CST-KO^ microbiota was constituted with species from *Proteobacteria*, *Patescibacteria*, *Desulfobacterota*, and *Coriobacteriia*. The increased abundance in those taxa in the WT ^FMT-CST-KO^ mice coincides with their higher abundance in the donor CST-KO, which showed a dominance of *Proteobacteria*, *Patescibacteria*, *Bacilli*, *Desulfobacterota*, and *Actinobacteriota*, further confirming the success of the FMT procedure.

Changes in microbial composition are often accompanied by metabolic changes, in particular, the production of short-chain fatty acids (SCFAs) (Tsukuda et al., 2021). Thus, levels of acetate, butyrate, and propionate, were measured in the cecum of donor and recipient mice of each genotype. Specifically, butyrate was significantly lower in the donor CST-KO mice compared to their WT counterparts as reported previously (**Figure 1F**) (González-Dávila et al., 2021). Levels of acetate and butyrate increased significantly in CST-KO^FMT-WT^ compared to CST-KO **(Figure 1F)**. In WT^FMT-CST-KO^ mice, no changes were visible compared to the WT control group. These observations are consistent with the observed significant increase in the butyrate and acetate-producing genus *Intestimonas* in the CST-KO^FMT-WT^ mice (Bui et al., 2015). As it is the only well-characterized and significantly changing taxon, this suggests that this specific genus is responsible for the strong increase in SCFA levels. Overall, the results imply that donor-specific taxa reliably colonized the recipients. Despite more prominent specific taxonomical changes in the WT ^FMT-CST-KO^ mice, FMT altered microbial richness and SCFAs production in CST-KO^FMT-WT^ but not in the WT^FMT- CST-KO^ recipient mice.

### Adoptive transfer of dysfunctional epithelial barrier and colonic fibrosis from CST-KO to WT mice

The gut microbiota has been strongly linked to intestinal homeostasis (El Aidy et al., 2012). Especially, in patients with IBD or colorectal cancer, an altered gut microbiota has been associated with a dysfunctional epithelial barrier and tissue inflammation, which, in turn, leads to sub-mucosal fibrosis (Franzosa et al., 2019; Halfvarson et al., 2017). Importantly, CST-KO mice have recently been shown to exhibit these disrupted intestinal mucosal processes, e.g. increased length and diameter of tight junctions, adherens junctions, and desmosomes, all coinciding with increased gut permeability (Muntjewerff, Tang, et al., 2021). Given the altered microbiota composition in CST-KO mice **(Figure 1;** (González-Dávila et al., 2021)), we hypothesized that the gut microbiota may play a causal role in the development of these features. To test our hypothesis, we employed whole-genome transcriptomic analysis on colonic tissue, as well as transmission electron microscopy (TEM) to examine the morphology of the intestinal colonic epithelium as well as the sub-mucosa of CST-KO and WT controls and the WT^FMT-CST-KO^ and CST-KO^FMT-WT^ recipient mice **(see methods and supplementary information for details)**.

The intestinal epithelium is regulated by a series of intercellular junctions between polarized cells: an apical tight junction (TJ) which guards paracellular permeability, the subjacent adherens junction (AJ), and desmosomes, both provide essential adhesive and mechanical properties that contribute to paracellular barrier functions (Buckley & Turner, 2018). On the transcriptome level, several genes involved in cell and TJ regulation and multiple TJ-markers such as Occludin (*Ocln*), MARVEL domain-containing protein 2 (*Marveld2*), Tight junction proteins (*Tjp1-3*) (Ma et al., 2004), as well as the desmosomal protein Desmoglein 2 (*Dsg2*), which is required for the maintenance of intestinal barrier function (Gross et al., 2018), and AJ genes coding for alpha-E-Catenin (*Ctnna1*), all showed substantial downregulation in WT^FMT-CST-KO^. In contrast, claudins (*Cldn1-6*) showed inconclusive expression profiles, as some were upregulated in WT^FMT-CST-KO^ but also CST-KO^FMT-WT^ (**Figure 2A**). The transcriptome data was further confirmed by ultrastructural examination of the gut epithelium using TEM, which revealed that the TJ, AJ, and desmosome morphology was different in CST-KO compared to WT, mostly exhibiting increased diameter for all three components, in line with previous findings (Muntjewerff, Tang, et al., 2021) **(Figure 2B)**. The fecal transplantations affected the morphology of the epithelial junctions of the recipient mice. TJs, AJs and desmosomes diameter appeared to be decreased in CST-KO^FMT-WT^, whereas in WT^FMT- CST-KO^ the diameter increased significantly compared to their donors. The TJs and AJs length was not affected in mice receiving FMT. However, desmosomes were significantly elongated in CST- KO^FMT-WT^ compared to CST-KO, while in WT^FMT-CST-KO^ there were no significant changes compared to WT. Collectively, the findings observed from the epithelial junction morphology indicated a phenotype transfer from their donor’s microbiota.

**Figure 2:**
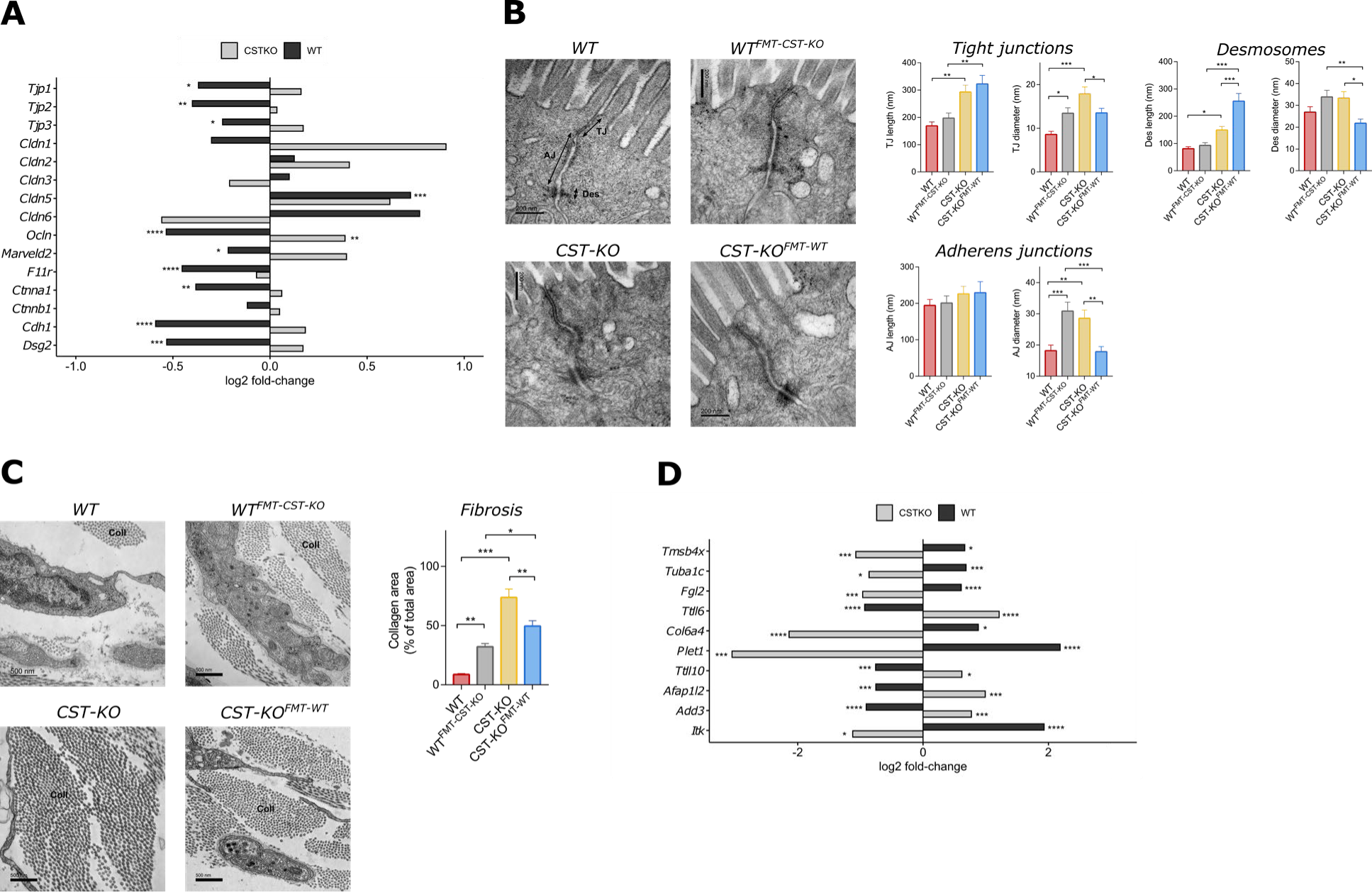
Adoptive transfer of colonic distortions from CST-KO to WT mice. **(A)** Barplot of genes involved in epithelial barrier regulation, showing log2 fold-changes for either CSTKO-KO vs CST-KO^FMT-WT^ or WT vs WT^FMT-CST-KO^. Log2 fold-changes and significance (padjusted < 0.05) of that change were determined by general differential expression analysis using *DESeq2* (see **Supplementray Excel Sheet 2**) **(B)** Representative transmission electron microscopy (TEM) micrographs showing epithelial barrier morphology for each group of mice. Arrows indicate the different parts: TJ – tight junction, AJ – adherens junction, Des – desmosome. The right side of the panel presents morphometric analyses results for TJ, AJ and Des, assessing length and diameter respectively; Bars show mean ± SEM **(C)** TEM micrographs showing the collagen fibers (cross-sectional view) for each group of mice. The images are representative for each group. Coll– collagen. The right side of the panel shows results of morphometric analysis showing fiber density as an assessment of fibrosis; Bars show mean ± SEM **(D)** Barplot of genes related the development of fibrosis, showing log2 fold-changes for either CSTKO-KO vs CST-KO^FMT-WT^ or WT vs WT^FMT-CST-KO^.

Moreover, the ultrastructural findings revealed by TEM demonstrated an excessive accumulation of the extracellular matrix component, collagen in the colonic tissue of WT^FMT-CST-KO^ and CST-KO mice compared to the CST-KO^FMT-WT^ and WT mice **(Figure 2C)**. Excessive accumulation of collagen is indicative of intestinal fibrosis, which is a characteristic of CST-KO mice (Muntjewerff, Tang, et al., 2021). Fibrosis results from tissue inflammation and has been associated with the upregulation of several immune-related genes, which are linked to the prognosis of IBD (Speca et al., 2012). Indeed, several genes involved in innate and adaptive immune response, cell adhesion, migration, proliferation, angiogenesis, skeletal development, and tissue wound repair were differentially regulated in the recipient groups compared to the donors **(Figure 2D)**. Altogether, the results infer that the CST-KO-associated microbiota may play a key role in the development of the altered tight junction regulation and fibrosis in CST-KO mice as inferred from the phenotype transfer after fecal microbiota transplantation in WT^FMT-CST-KO^, and that FMT from WT donors reversed, to a great extent, this distortion as shown in CST-KO^FMT-WT^.

### Core regulatory network that governs the transcriptional changes and their association with the transplanted microbiota

In order to gain a more mechanistic understanding on how the transferred microbiota interfaces with the host and which pathways are involved to express a specific phenotype, we performed a comprehensive analysis of the obtained transcriptomic data. A total of 5233 genes were differentially expressed in the colonic tissue of WT^FMT-CST-KO^ compared to the WT mice, with 2712 (log2FC >2: 765) of these genes being upregulated and 2521 (log2FC >2: 295) downregulated **(Figure 3A)**. In CST-KO^FMT-WT^, a total of 4273 genes were found to be differentially expressed in the colonic tissue compared to the CST-KO mice, with 1987 (log2FC >2: 629) of these genes being upregulated and 2286 (log2FC >2: 916) downregulated **(Figure 3A)**. Principal component analysis (PCA) performed on normalized gene expression showed a similar convergent clustering of the recipient mice compared to the donors **(Figure 3B)**, in line with the microbiota clustering **(Figure 1C, Supplementary Figure 1).**

**Figure 3:**
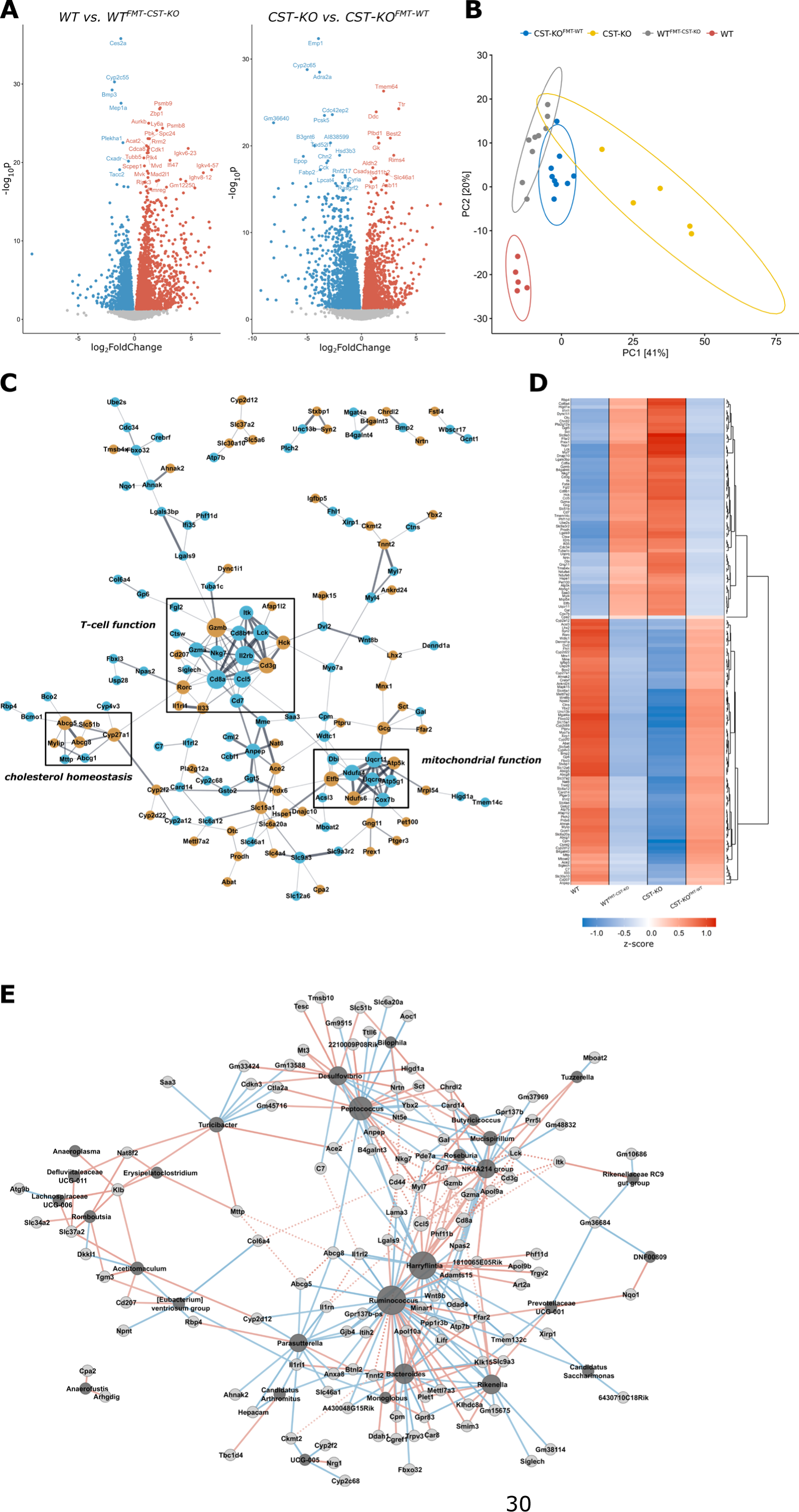
Core regulatory network that governs the transcriptional changes and their association with the transplanted microbiota. **(A)** Volcano plots of differentially expressed genes comparing either WT vs WT^FMT-CST-KO^ or CST-KO vs CST-KO^FMT-WT^. Log2 fold-changes and significance (padjusted < 0.05) of that change were determined by general differential expression analysis using *DESeq2* (see **Supplementray Excel Sheet 2**). Genes significantly upregulated in the FMT groups are highlighted in red, while significantly downregulated genes are colored in blue. **(B)** Principal component analysis (PCA) of normalized gene expression data revels distinct clustering between all of the groups. Ellipses represent normal data ellipses produced by methods from the R package ggplot2. **(C)** Network representation of genes regulated in opposite direction in the recipient groups (WT^FMT-CST-KO^ and CST-KO^FMT-WT^ compared to their donors: i.e., the same gene is upregulated in one recipient group but downregulated in the other recipient group; log2 fold- change >0.58, padjusted < 0.05). Connections are inferred from the STRING database (combined score >0.4), while edge width corresponds to interaction strength. Node size is determined by its degree. Brown colour indicates involvement in IBD determined by mining from the TaMMA meta- transcriptome catalogue (Massimino et al., 2021) **(D)** Heatmap of all core genes (including the ones shown in (C)) clustered by their group/gene z-score normalized expression, illustrating the dualistic effect of FMT for those genes in each genotype. **(E)** Network representation of correlation analysis (Pearson, padjusted < 0.05) between the abundance of microbial taxa and the expression of certain genes presented in (D). The network is only representative for WT. Light gray – Gene, Dark gray – Microbial taxon, Blue – negative correlation, Red – positive correlation; Solid lines refer to direct correlation, while dotted lines depict interactions inferred from the STRING database.

A comprehensive functional analysis of altered biological processes was carried out between the recipient and donor mice and revealed alterations in immunological processes, including immunoglobulin production, B-cell activation and adaptive immunity regulation to be upregulated in WT^FMT-CST-KO^ compared to the WT control. In contrast, changes in CST-KO^FMT-WT^ were characterized by downregulation of lipid metabolism and ion transport compared to the CST-KO mice (Detailed analysis can be found in (**Supplementary Figures 2 and 3**). Overall, the transcriptome data revealed a significant impact of the FMT on the transcriptome level in both CST-KO and WT mice. From the differential expression analysis, we hypothesized that there may be a core set of regulatory genes that could serve as transcriptional signatures for the altered mucosal homeostasis upon adoptive microbiota transfer between CST-KO and WT mice in the colon. Therefore, all DEGs with an opposite differential regulation in the recipient groups (WT ^FMT-CST-KO^ and CST-KO^FMT-WT^ compared to their donors: i.e., the same gene is upregulated in one recipient group but downregulated in the other recipient group; (log2-fold change >0.58, p adjusted <0.05; see **(Supplementary Excel Sheet 2)**, were mined to search for potential transcriptional signatures **(see Methods for detailed description)**. A total of 292 genes were found to be commonly expressed in both the CST-KO ^FMT-WT^ and WT ^FMT-CST-KO^, but in opposite directions. The associations among the proteins translated by the commonly identified DEGs in the recipient groups in comparison to their controls were assessed and interactions among the query proteins were further visualized. This resulted in a network that comprised 143 nodes and 233 edges of protein-protein interactions after excluding unconnected nodes or clusters of only two genes **(Figure 3C)**. The resulting network exemplified the strong impact of adoptive microbiota transfer between CST-KO and WT mice in the colon on both immune and metabolic gene expression throughout the colonic mucosa and encompassed several core regulatory genes that are known to control the induction of metabolic and immune responses. Cluster analysis was used to identify closely interlinked regions from the network of proteins **(Figure 4C)**. The top 3 clusters were found to be highly significant and included the most nodes out of the ones reported (**Supplementary Table 1**). Using functional enrichment of the clustered proteins, potential biological processes were assigned to each cluster. Each cluster was related to a separate process, including genes annotated to belong to T-cell function, energy metabolism (mitochondrial functions), and cholesterol homeostasis.

Genes from the cluster involved in T-cell function exclusively showed higher expression in WT^FMT- CST-KO^ and CST-KO donors, indicating potential activation of T-cell-related immunological pathways by CST-KO transplanted microbiota. A similar expression pattern was also found for the genes in the energy metabolism cluster. Intriguingly, the genes present in the cholesterol homeostasis cluster were found to be enriched (except *Slc51b*) in CST-KO^FMT-WT^ and WT donors, indicating a simulative effect of the WT-transferred microbiota (**Figure 3D**). To further assess whether FMT was able to restore the expression levels of the genes identified in the core regulatory network to their original expression levels in the recipient groups, i.e., showing no significant differential expression anymore, we performed a comparative analysis between the CST-KO^FMT-WT^ vs WT and WT^FMT-CST-KO^ vs CST-KO groups. Among all genes in the identified transcriptome signatures, 234 were found to be restored, while 134 were also present in the core network **(Supplementary Excel Sheet 2)**. Considering the high fractions of the identified genes, the data highlight the powerful efficacy of the FMT to restoring the transcriptional patterns.

Next, possible associations between the colonic microbiota, which was transferred from the donor to the recipient genotypes **(Figure 1),** and the core regulatory genes **(Figure 3D)** were explored using pairwise Pearson correlations. The correlation analysis revealed that 41% of the core differentially expressed genes were associated with at least one microbial taxon, which were detected to be significantly differentially abundant, only in the WT^FMT-CST-KO^ recipient group **(Supplementary Excel Sheet 2)**. The associations between core network gene expression and taxon abundance, as well as interaction among the query proteins, were further visualized (**Figure 3E**). Five main clusters were identified, among them, one was found to be less connected to the others, mostly containing low-degree taxon nodes. *Ruminococcus*, *Harryflintia,* and *Peptococcus* had the most correlations among the taxa (average of 29 correlations), while *Klb*, *Klk15*, and *Myl7* were the most connected genes (6, 5, and 6 edges, respectively). Genes from the core T-cell cluster and generally immune-related genes connected to *Peptococcus, Harryflintia, NK4214 group, Desulfovibrio,* and *Ruminococcus*. Genes from the energy metabolism cluster did not show any apparent significant correlations to any of the taxa, while genes involved in cholesterol homeostasis (including core gene cluster 3), such as *Mttp*, *Acbg5/8*, and *Slc51b* were correlated with *Turicibacte*r, *Peptococcus, Parasutterella*, *Desulfovibrio, and Bilophila*.

As CST is proposed to play a crucial role in IBD, we investigated which genes from our core regulatory network are involved in this disease. Intriguingly, among the identified tissue transcriptome signatures, the human counterparts of 108 genes (62 genes in the identified core network) have a known role IBD determined by matching them to the TaMMA IBD transcriptomics catalog (Massimino et al., 2021), and 36 of which are linked to bacteria. These findings further support the biological relevance of the identified transcriptional signatures for mucosal control of homeostasis along the gut and the role of altered microbiota composition in CST-KO mice as a cause in differential expression of genes involved in mucosal immune and metabolic processes.

## Discussion

The present findings demonstrates that CST-KO-associated alterations in the gut microbiota are sufficient to disrupt colonic homeostasis in healthy mice. Similarly, the transfer of the gut microbiota of healthy mice restored the distorted colonic function in CST-KO mice. Specifically, transplantation of the perturbed microbiota signature from CST-KO mice to microbiota-depleted WT mice induced the development of several colonic dysfunctional features of the CST-KO phenotype on the gene and tissue levels. CST-KO mice are associated with an altered gut microbiota composition, and richness **(Figure 1** and González-Dávila et al. 2021). A major parallel between the CST-KO microbiota profiles and the WT mice that received the FMT from the CST-KO (WT^FMT-CST-KO^) encompassed a reduction of Clostridia and *Akkermansia*, which has been previously linked to metabolic disorders and insulin resistance (Schneeberger et al., 2015; Zhou et al., 2021), a CST-KO related phenotype (Bourebaba et al., 2021) as well as a prominent increase in the *Proteobacteria* population, all of which have been found previously to be indicative for active IBD states (Earley et al., 2019; Glassner et al., 2020). In contrast, CST-KO that received the FMT from the WT mice (CST-KO^FMT-WT^) encompassed an increase in richness and a notable reduction of *Staphylococcus*, as well as an increase in the butyrate-producing *Intestinimonas* **(Figure 1).** In fact, reduced levels of butyrate have been strongly linked to IBD as well as metabolic disorders (De Preter et al., 2011; Huda-Faujan et al., 2010).

Detailed transcriptome analysis allowed the identification of transcriptional signatures of genes commonly differentially regulated in recipient mice in opposite directions **(Figure 3C, D)**. These transcriptional signatures included the correlative expression of metabolism-related genes and immune-related genes. Most of these genes showed full restoration of their transcriptional levels when compared to their opposite controls. Among the identified transcriptional signatures, were several genes known to be involved in fibrosis; which was indeed confirmed by TEM, where an excessive accumulation of the extracellular matrix component, collagen was observed in the colonic tissue of WT^FMT-CST-KO^ and CST-KO mice compared to the CST-KO^FMT-WT^ and WT mice **(Figure 2C)**. Furthermore, the identified signatures comprised genes of which the human orthologues are dysregulated in IBD. Based on genes identified in large-scale transcriptomic meta-analysis (TaMMA), at least 37% of the found genes were associated with inflammatory bowel disease.

Our transcriptome and transmission electron microscopy data demonstrated that transferring the gut microbiota from CST-KO mice to WT mice with a depleted gut microbiota could induce the development of some of the features of the dysfunctional colon, such as distorted barrier integrity and fibrosis resulting in a physiological profile similar to CST-KO **(Figure 2)**. Further, GO enrichment and pathway analysis consistently identified “immune response” as the main umbrella category affected in response to FMT from CST-KO mice. CST-KO mice exhibited an increased expression in pro-inflammatory genes, namely *Infg, Itgam, Itgax, Cxcl1, Il12b,* and *Nos2* which, were also increased in WT^FMT-CST-KO^ **(Supplementary Figure 3 and Supplementary Excel Sheet 2)**. The results are in line with the recent findings showing that CST-KO mice have infiltration of macrophages and CD4^+^ T-cells, and higher gene and protein expression levels of pro-inflammatory molecules in the gut, indicative for IBD-like states (Muntjewerff, Tang, et al., 2021). The same study (Muntjewerff, Tang, et al., 2021) reported that CST-KO mice have increased gut permeability, altered tight junctions morphometry, which is in agreement with the decreased gene expression of several TJ-markers such as *Clnd5, Dsg2, Ctnna1, Tjp2, Tpj3, Ocln, Marveld2* in CST-KO **(Figure 2A)** as well as their distorted TJ, AJ, and desmosome morphology **(Figure 2B)**, compared to WT, all of which (except for *Cldn5*) were retored in WT^FMT-CST-KO^ mice decreased the gene expression in the same TJ-markers.

In contrast to the induced immune response in WT mice that received FMT from CST-KO, CST- KO^FMT-WT^ mice displayed a downregulation in several cell processes involved in lipid metabolism, suggesting that metabolic reorientation of colonic tissue from an oxidative energy supply had occurred. Notably, CST is known to be involved in body weight regulation through its effects on lipolysis and fatty acid oxidation, as well as its beneficial effects in mice with diet-induced obesity (Bandyopadhyay et al., 2012; Ying et al., 2018a), and is correlated with plasma HDL-cholesterol levels (Durakoğlugil et al., 2015). However, such changes are usually of systemic nature, affecting the expression of genes in adipose or liver tissue. In the intestinal milieu, *Mttp*, *Abcg5* and *Abcg8*, which have been attributed to cholesterol/triglyceride uptake are elements of the core regulatory network that regulate the transcriptional changes and their association with the transplanted microbiota **(Figure 3C-E)**, altogether indicating strong cross-correlation in their expression (Lally et al., 2006). Intestine-specific ablation mutants of *Mttp* showed a decrease in cholesterol transport from the intestine to the plasma and a subsequent reduction in the plasma cholesterol and triglyceride levels, but an accumulation of triglycerides in the intestine (Iqbal et al., 2013). Alterations in lipid metabolism and elevated immune responses are interrelated, and the gut microbiota was shown to play a key role in this connection (El Aidy et al., 2013). Patients suffering from IBD were also found to have major alterations in lipid metabolic processes both in intestinal tissue as well as systemically (Heimerl et al., 2006; Scoville et al., 2018). More generally immune responses due to pathogen infection have been linked to abnormal energy metabolism in chicken (Wang et al., 2020), further confirming a strong interplay between metabolic reorientations in the intestine and the restoration of inflammatory responses. Gut microbiota metabolises lipids in the intestinal lumen, consequently affecting the host (lipid) metabolic profile (Lamichhane et al., 2021). Specifically *Turicibacter*, which, in our data, was decreased in WT^FMT-CSTKO^ compared to WT, and showed a strong positive correlation with the core gene network, in particular with *Mttp* from the cholesterol homeostasis gene cluster **(Figure 3E),** has been associated with the modulation of lipid metabolism in mice (Fung et al., 2019). This, in conjunction with the significant increase in butyrate in the CST-KO mice that received the WT FMT (**Figure 1B**), suggests that FMT caused a change in the microbiota metabolic output, with a consequent shift in host metabolism and epithelial barrier restoration.

Overall, our data suggest that the gut microbiota may play a causal role in the complex mechanisms underlying the development of diseases related to altered levels of CST, such as IBD and metabolic diseases. The profile of physiological and gene alterations in the colon following FMT may represent a novel paradigm in intestinal pharmacology to investigate potential microbiota-associated disorders. The identified transcriptome signatures included several genes of which the human orthologues are IBD and metabolic disease-associated genes that have also been discovered in large-scale transcriptome studies, suggesting their relevance for the mucosal control of homeostasis, and supporting their importance in the dysregulation of immune- and metabolic-associated pathways in patients with altered levels of CST. Findings from this study may advance the concept that targeting the gut microbiota could be a viable therapeutic strategy for a novel development of diseases associated with altered levels of CST, therefore may augment prevention strategies in these diseases.

## Materials and Methods

### Animals

All studies with mice were approved by the University of California San Diego (UCSD) and Veteran Affairs San Diego Healthcare System (VASDHS) Institutional Animal Care and Use Committees for the Mahata laboratory (UCSD: #S00048M; VA: #A13-002) and were performed in San Diego, California in adherence to the NIH Guide for the Care and Use of Laboratory Animals. For the FMT experiment twelve male adult C57BL/6 J mice (age 20 weeks) were purchased from Jackson Laboratory (Bar Harbor, ME) and were acclimatized for 3 weeks before experimentation. Additionally, twelve CST-KO mice generated in the Mahata laboratory were used for this experiment. CST-KO mice have a deletion in the 63 bp CST domain from Exon VII of the *Chga* gene ^27^ in C57BL/6 background. For transcriptome analysis, an additional 5 and 7 age and sex matched mice from WT and CST-KO respectively, served as controls. Mice were housed in 4 to 5 animals per cage and had free access to water and food (Normal Chow Diet, LabDiet 5001) in temperature and humidity-controlled rooms with a 12-hr light/dark cycle.

### Fecal microbiota transplantation

After 3 weeks of acclimatization period, FMT was performed reciprocally: CST-KO animals received a fecal transplant from WT donors and WT animals received a fecal transplant from CST-KO donors. To avoid the adverse effects of antibiotic treatment on the intestinal transcriptome and physiology (Allegretti et al., 2018), bowel cleansing was performed with polyethylene glycol (PEG), an osmotic laxative agent, as previously described (Wrzosek et al., 2018). In brief, CST-KO and WT mice (n= 12) were fasted 4 hr before bowel cleansing, with free access to water. Mice received four consecutive bowel cleansings of 425 g/L PEG4000 (Sigma Aldrich) by oral-gastric gavage. Each round of bowel cleansing was performed at 20 min intervals. Due to the laxative effect of PEG, the animals were moved constantly to sterile clean cages without bedding to avoid coprophagy and re-inoculation of the former microbiota. To verify the success of the cleansing procedure, sample mice (n= 2) were chosen randomly, euthanized and the total gut was checked to detect luminal content and it was compared to the intestinal content of a control mouse treated with 0.9% saline **(Supplementary Figure 4)**.

Bowel cleansing was only performed on day 1 Five hours later, mice received the first fecal transplant (200 μl fecal suspension) by oral-gastric gavage. In total, mice received 3 fecal suspensions, one per day. Fecal suspension for FMT was prepared as following one day before FMT, a pool of feces from 10 mice/group were collected freshly, frozen immediately, and stored at - 80°C. The fecal suspension (1:10 w/v) was prepared each day of FMT using the frozen pool of feces and sterile PBS. The solution was vortexed gently for 10 min and centrifuged at 800 rpm for 3 minutes. Only the supernatant was used for FMT oral gavage. After the first FMT, mice had normal access to food, water, and housing conditions. Mice were sacrificed 14 days after recolonization.

### DNA isolation and 16S sequencing

DNA extraction was performed on fecal samples that were collected before and post-FMT, using a phenol-chloroform-isoamyl alcohol procedure (Santella, 2006). Briefly, the pellet was resuspended in 1 ml of lysis buffer (940µl TE buffer, 50 µl SDS 10% and 10 µl Proteinase K 20 mg/ml) in a 2 ml screw cap microtube containing a mix of zirconium and glass beads. Then, samples were incubated at 58°C for 1 hour, and 150 µl buffered phenol (Invitrogen, 15513-047) was added. In each step, samples were vigorously mixed using a vortex. To support lysis, samples were homogenized 3 × 30 sec with 1-min intervals on ice in a mini bead-beater (Biospec, Bartlesville, USA). This was followed by the addition of 150 µl chloroform/isoamyl alcohol (24:1) and centrifuged at 16,000x *g* for 10 min at 4°C. The upper layer, which contains the DNA, was carefully transferred to a clean tube and 300 µl of phenol/chloroform/isoamyl alcohol [25:24:1] was added. Again, the upper layer was transferred to a fresh tube and 300 µl of chloroform/isoamyl alcohol [24:1] was added. To precipitate the DNA, the upper layer was transferred to a new tube and 1 volume of absolute isopropanol and 1/10 volume of 3M sodium acetate was added. Samples were incubated overnight at -20°C. To get the DNA pellet, the samples were centrifuged at 16,000x *g* for 20 min at 4°C. The supernatant was removed and 700 µl of 70% ethanol was added to remove remaining salts from the pellet. Ethanol was removed and the DNA pellet was air-dried for 30 min before resuspension of the pellet in 100 µl TE buffer.

Sequencing of the V3-V4 region of the bacterial 16S gene was carried out by Novogene Co. Ltd. Briefly, for sequencing library preparation, raw DNA extracts were diluted to 1ng/µl in sterile water, and amplicons were generated by PCR (primers 341F and 806R) using a Phusion® High-Fidelity PCR Master Mix (New England Biolabs). Amplification product quality was assessed by gel electrophoreses and samples were pooled in equimolar ratios. Libraries were generated with a NEBNext® UltraTM DNA Library Prep Kit for Illumina and sequencing was carried out on an Illumina 250bp paired-end platform. Initial processing of reads involved trimming of adapters and primers using a Novogene in-house pipeline (Novogene Co. Ltd, Cambridge, UK).

### Microbiota analysis

Paired-end sequencing reads were filtered, denoised, merged, and classified with the *dada2* package in the statistical programming language R while processing forward and reverse reads separately until merging (Callahan et al. 2016; R Core Team 2019). Briefly, reads were truncated to 220bp and low-quality reads were filtered followed by dereplication. Error models were learned while manually enforcing the monotonicity of the error function. Reads were denoised and merged with a minimal overlap of 12bp, while non-merging reads were concatenated. Singletons were removed before performing bimera removal, followed by read classification using SILVA (V138) as a taxonomical reference database.

For downstream analysis, the *phyloseq* and *microbiome* packages to determine richness and alpha diversity were assessed via observed amplicon sequence variants (ASVs), chao1, Shannon’s *H,* and inverted Simpson’s index (Lahti and Shetty n.d.; McMurdie and Holmes 2013). All ASVs were collapsed on the genus level and cumulative sum scaling was applied using *metagenomeSeq* (Paulson et al. 2013). The resulting genus abundance table served as input for different types of ordinations such as principal component analysis (PCA) or redundancy analysis (RDA). To test for differences between all clusters, PERMANOVA was applied using the *adonis* function from the *vegan* package (Oksanen et al. 2019). Differential abundance between before and post-FMT groups was assessed by paired Wilcoxon Rank Sum test followed by FDR correction using p < 0.05 as a significance threshold. Additionally, LefSe analysis (Linear discriminant analysis Effect Size Segata et al. 2011) was carried out, using standard parameters.

### Determination of short-chain fatty acids in cecal samples

Cecal samples (100 mg) were suspended in 1 mL of saturated NaCl (36%) solution. An internal standard (50 μL of 10.7 µM 2-ethylbutyric acid in MQ water) was added and the samples were homogenized using glass beads. After the addition of 150 µL H2SO4 96%, SCFAs were extracted with 3 ml of ether. The ether layer was collected and dried with Na2SO4 (150 mg). The supernatant (0.5 µL) was analyzed using gas chromatography with flame ionization detection (Agilent, Santa Clara, California, USA). The system was equipped with a DB FFAP analytical column (30m x 0.53 mm ID, 1.0 µm; Agilent) and helium GC grade (5.6) was used as carrier gas with a constant flow of 4.2 ml/min. The initial oven temperature was held at 100 °C for 3 min, ramped with 4 °C/min to 140 °C (isothermal for 5 min) and further with 40 °C/min to 235 °C (isothermal for 15 min). Graphs and statistical analysis were performed with GraphPad Prism, using unpaired t-tests. Significance is indicated in the figure legend.

### Transmission Electron Microscopy (TEM) and morphometric analysis in mice colon

To displace blood and wash tissues before fixation, mice were deeply anesthetized and were cannulated through the apex of the heart and perfused with a pre-warmed (37 C) calcium and magnesium buffer with 10 mM KCl for 3 min followed by perfusion with freshly prepared pre-warmed (37 C) fixative containing 2.5% glutaraldehyde, 2% paraformaldehyde in 0.15 M cacodylate buffer for 3 min as described previously (Pasqua et al., 2016). The mouse colon was dissected. The fixation, embedding, sectioning, and staining of the mouse colon were performed as described also by . Grids were viewed using a JEOL JEM1400-plus TEM (JEOL, Peabody, MA) and photographed using a Gatan OneView digital camera with 4×4k resolution (Gatan, Pleasanton, CA). Micrographs were randomly taken from 3 mice per group (8 to 10 photographs per mouse with a total of 25 to 28 photographs) and the morphometry of tight junctions and fibrosis were performed as described previously (Muntjewerff, Tang, et al., 2021; Pasqua et al., 2016). Briefly, the line segment tool in NIH ImageJ was used to measure the lengths and perpendicular widths (diameter) of TJ, AJ, and desmosomes. The free-hand tool in NIH ImageJ was used to manually trace around the area occupied by the collagen fibers. For determination of the collagen area, the sum of the collagen area in a randomly-chosen photograph was divided by the total area of that photograph and multiplied by 100.

### RNA extraction from colonic samples

Samples were collected and submerged in RNA later (Qiagen) to avoid RNA degradation and stored at -80 °C. A hybrid protocol using TRIzol (Invitrogen) and RNeasy Mini Kit (Qiagen) was used to obtain high-quality RNA. Briefly, 10-20 mg of tissue was submerged in 1 ml ice-cold TRIzol in a 2 ml screw cap microtube containing 3 mm glass beads. To perform lysis, samples were homogenized 3 × 30 sec with 1-min intervals on ice in a mini bead-beater (Biospec, Bartlesville, USA). The sample was centrifuged at 12,500x *g* for 15 min at 4 °C. Then, it was transferred to a clean tube where 200 μl of chloroform was added and mixed with a vortexer. After centrifuged at 12,500x *g* for 15 min at 4 °C, the upper clear layer was collected in a fresh tube and two volumes of 100% ethanol were added and mixed gently. Immediately, the mixture was transferred to an RNAeasy mini kit column and centrifuged for 30 sec at room temperature. To wash the RNA, two times 500 μl RPE solution was added, and RNA was eluted in a tube with RNAse-free water. RNA quality was assessed by gel electrophoresis.

### RNA sequencing in colon samples

RNA library was assembled using NEBNext Poly(A) mRNA Magnetic Isolation Module (E7490) and NEBNext Ultra II RNA Library Prep Kit for Illumina (E7770, New England Biolabs) as per manufacturer instructions. Single-end sequencing was performed using a NextSeq 500 machine (Illumina; up to 75 cycles). The generated data were subsequently demultiplexed using sample-specific barcodes and changed into fastq files using bcl2fastq (Illumina; version 1.8.4). The quality of the data was assessed using FastQC (v0.11.8) (Andrews, 2010). Low quality bases and (parts of) adapter sequences were removed with Cutadapt (v1.12; settings: q=15, O=5, e=0.1, m=36) (M. Martin, 2011). Sequenced poly-A tails were removed as well, by using a poly-A and a poly-T sequence as adapter sequences (A{100} and T{100}). Reads shorter than 36 bases were discarded. The trimmed fragment sequences were subsequently aligned to the mouse reference genome (GRCm39; From Ensembl; release 103) and the number of reads per gene were determined with the use of STAR ((Dobin et al., 2013); v2.7.8a; settings: -- outSAMstrandField=intronMotif, --quantMode=GeneCounts, --outFilterMultimapNmax=1). Duplicate reads were marked with samtools markdup (v1.9; using htslib 1.9) and the extent of PCR amplification (PCR artefacts) was assessed with the use of the R package dupRadar (v.16.0) (H. Li et al., 2009; Sayols et al., 2016). There was no indication of PCR artifacts for any of the samples (Intercept: 0.006 – 0.013; Slope: 4.50 – 5.69; webpage of dupRadar was used as a guideline; https://bioconductor.org/packages/release/bioc/vignettes/dupRadar/inst/doc/dupRadar.html).

### Differential gene expression and gene ontology analysis

The principal component analysis was performed in R (v3.6.3) using the R package *DESeq2* (v1.26.0) (Love et al., 2014; R Core Team, 2019). To visualize the overall effect of experimental covariates as well as batch effects (function: *plotPCA*). Differential gene expression analyses were performed with the same R package using default settings (Negative Binomial GLM fitting and Wald statistics), following standard normalization procedures.

The function enrichGO of the Bioconductor R package clusterProfiler (v3.14.3) was used to test whether certain gene ontology (GO) categories were enriched among the detected genes (settings: OrgDb=org.Mm.eg.db [v3.10.0], keyType=ENSEMBL, universe=[all genes with an adjusted p-value; padj is not NA], qvalueCutoff=0.05, minGSSize=1, maxGSSize=100,000), while only considering the bioprocess aspect of GO (Carlson, 2019; Yu et al., 2012).

The expected number of genes for each category (see barplots) were calculated with the gene ratio (GeneRatio) and background ratio (bgRatio) information from the output of the enrichGO function (background ratio times total number of genes with GO annotation that were differentially expressed).

### Differential gene expression analysis with GSEA

Normalized expression levels were subjected to Gene Set Enrichment Analysis (GSEA) using default parameters (Subramanian et al., 2005). GO bioprocess was used as gene set category and enrichment results were visualized using the EnrichmentMap plugin (v3.3.3) of Cytoscape (v3.9), by considering all enriched categories below a nominal p-value of 0.05 (Merico et al., 2010). Clusters were grouped and annotated using the Autoannotate plugin (v1.2) with default parameters.

### Construction of core regulatory network

DEGs (padjusted<0.05) were subset to contain only genes which exhibited an opposite foldchange upon FMT, being greater than +-0.58 log2FC. Associations among the proteins translated by the commonly identified DEGs were assessed using the StringApp plugin (v1.7) from Cytoscape (v3.9) (Doncheva et al., 2019; Shannon et al., 2003). The minimum required interaction score was set to 0.4 and interactions among the query proteins were further visualized. Single nodes and two node clusters were removed and clusters were identified with Cytocluster ClusterONE (v.1.0) (M. Li et al., 2017) with default settings. Subsequently, functional enrichment using StringApp was performed on the resulting clusters. Visualisation was carried out in Cytoscape.

### Correlation analysis of gene expression levels and microbial taxon abundance

Genes were subset to only contain genes which change in a significant manner upon FMT. Pairwise Pearson correlations were computed between normalised expression levels (DSeq2) for each gene against normalised abundance values (CSS) of each microbial taxon using the *psych* R package. This was done separately per genotype only on FMT timepoints. Only significant correlations were kept for further analysis. To overcome the fact of unpaired samples for the control groups, gene and taxon log2 fold-changes (log2FC) were used to discard any false positive correlations. When the sign of the correlation was positive the sign of the gene and taxon log2FC were required to be equal, contrary if the correlation sign was negative the signs of the log2FCs had to be different. To further refine the analysis correlations were subset to contain only genes which change in an opposite fashion, eg. positive log2FC in WT and negative log2FC in CSTKO, as well as an log2FC greater than 0.58. We performed graph analysis and visualisation in Cytoscape v3.9 also employing the StringApp plugin.

### Statistical analysis

All statistical tests were performed using GraphPad Prism 7 or the statistical programming language R. Specific statistical tests are indicated in the text or figure legends. Normality was assessed by either D’Augustino-Pearson omnibus normality test or Shapiro-Wilk normality test. If normality was met a t-test was chosen, otherwise non-parametric Mann-Whitney was used. Outliers were assessed with GraphPads ROUT method (Q=1).

### Availability of data and materials

All data generated or analyzed during this study are included in this published article and its supplementary information files. Whole transcriptome and 16S rRNA gene amplicon sequence data were deposited under BioProject numbers PRJNA800626 and PRJNA741992.

## Authors’ Contributions

P.G-D., M.S, S.E.A, and S.K.M conceived and designed the study. P.G-D., A.D., M.S., B.D, and S.K.M. performed the experiments, and P.G-D., M.S, R.W, B.D., S.K.M., and S.E.A analyzed the data. P.G-D, M.S, S.E.A. wrote the original manuscript that was reviewed by A.D, R.W., B.D., K.V., and S.K.M. Funding for these studies was acquired by S.E.A. and S.K.M. All authors read and approved the final manuscript.

## Acknowledgment

P.G-D thanks the National Council of Science and Technology in Mexico (CONACyt) for the Ph.D. grant assigned to CVU 690069. We thank the UMCG/ERIBA Research Sequencing facility for help with RNA sequencing. We further thank Dr. Danny Incarnato of Department of Molecular Genetics, University of Groningen, the Netherlands, for his guidance in the RNA-Library preparation; Dr. Greet Vandermeulen of Department of chronic diseases and metabolism, Faculty of Medicine, KU Leuven, Belgium for the help with SCFAs analysis.

