## Supplementary information for "Gut microbiota transplantation drives the adoptive transfer of colonic genotype-phenotype characteristics between mice lacking catestatin and their wild type counterparts"

### Supplementary Results

#### CST-KO animals show distinct transcriptional patterns compared to their WT counterparts

Transcriptome analysis was performed using *DeSeq2,* we detected a total of 3690 differentially expressed genes (DEGs) in CST-KO mice, 2195 were upregulated and 1765 were downregulated ($p_{adj} < 0.05$) compared to WT counterparts. To complement our analysis, we further employed the overrepresentation analysis (OA) with ClusterProfiler tool on the previously determined DEGs as an alternate method to determine significantly changing GO biological processes in CST-KO mice compared to WT **(Supplementary Figure 2, A-C)**. This analysis identified differentially expressed genes related to regulation of hormone metabolic processes, lipid metabolism, molecular transport and peptide secretion.

In parallel, we employed Gene Set Enrichment Analysis (GSEA) to search for groups of genes that are significantly enriched in either genotype (Subramanian et al., 2005). For the GSEA, we used again the Gene Ontology biological processes database consisting of 4055 gene sets out of which 26, and 35 gene sets were significantly enriched in CST-KO and WT respectively (nominal p-value < 0.05). GSEA identified enriched GO biological processes involved in catecholamine metabolic processes especially for norepinephrine uptake and secretion, maintenance of the gastrointestinal epithelium, nervous system development, T-cell mediated immunity and regulation in leukocyte chemotaxis **(Supplementary Figure 2, D-G)**. Collectively, the transcriptome data is in agreement with the known role of catestatin as a nicotinic cholinergic antagonist that affects the catecholamine release (Mahata et al., 1997), regulator of the gut epithelium barrier (Eissa et al., 2018; Muntjewerff et al., 2021; Rabbi et al., 2017), regulator of metabolic and immune homeostasis (Muntjewerff, Christoffersson, Mahata, & van den Bogaart, 2022; Muntjewerff, Dunkel, Nicolasen, & Mahata, 2018), anti-obesogenic, and regulator of lipid metabolism (Ying et al., 2018).

#### Enriched biological functions and pathways modulated in response to the adoptive microbiota transfer in CST-KO$^{FMT-WT}$ and WT$^{FMT-CST-KO}$.

To comprehensively describe which biological processes are altered in CST-KO$^{FMT-WT}$ and WT$^{FMT-CST-KO}$ compared to their controls, we employed GSEA on biological process gene ontology (GO) level. A total of 120 gene sets were enriched in CST-KO$^{FMT-WT}$ compared to the CST-KO mice (nominal p-value < 0.05), while 275 gene sets were enriched in the WT $^{FMT-CST-KO}$ compared to WT mice **(Supplementary Excel Sheet 2)**. To complement our analysis, we further used overrepresentation analysis (OA) with the ClusterProfiler tool on the previously determined DEGs as an alternate method to determine significantly changing biological pathways **(Supplementary Figure 3).** Collectively, both analyses identified several clusters of gene sets involved in immunological processes, including immunoglobulin production, B-cell activation and adaptive immunity regulation; regulation of innate immunity, immune system activation and MHC antigen processing, to be upregulated in WT $^{FMT-CST-KO}$ compared to the WT control. Additionally, GSEA identified several clusters of gene sets involved in upregulation of chromosome separation, chromatin remodelling, DNA replication and cell cycle-related gene sets in WT $^{FMT-CST-KO}$ compared to the WT control. OA additionally identified a large downregulated cluster related to cell regulatory functions in WT $^{FMT-CST-KO}$. Remarkably, changes in CST-KO$^{FMT-WT}$ were less prominent compared to the WT$^{FMT-CST-KO}$. Moreover, the CST-KO$^{FMT-WT}$ group displayed fewer upregulated genes but greater number of downregulated genes compared to the WT$^{FMT-CST-KO}$. GSEA identified a few clusters of gene sets involved in DNA repair damage, pyrimidine metabolic processes, glycolipid metabolism, blood pressure regulation, and vitamin (folate) metabolism, to be upregulated in CST-KO$^{FMT-WT}$ compared to CST-KO group. These results were supported by OA, which additionally showed upregulation in gene sets involved in lipid metabolism.

### Supplementary Figures

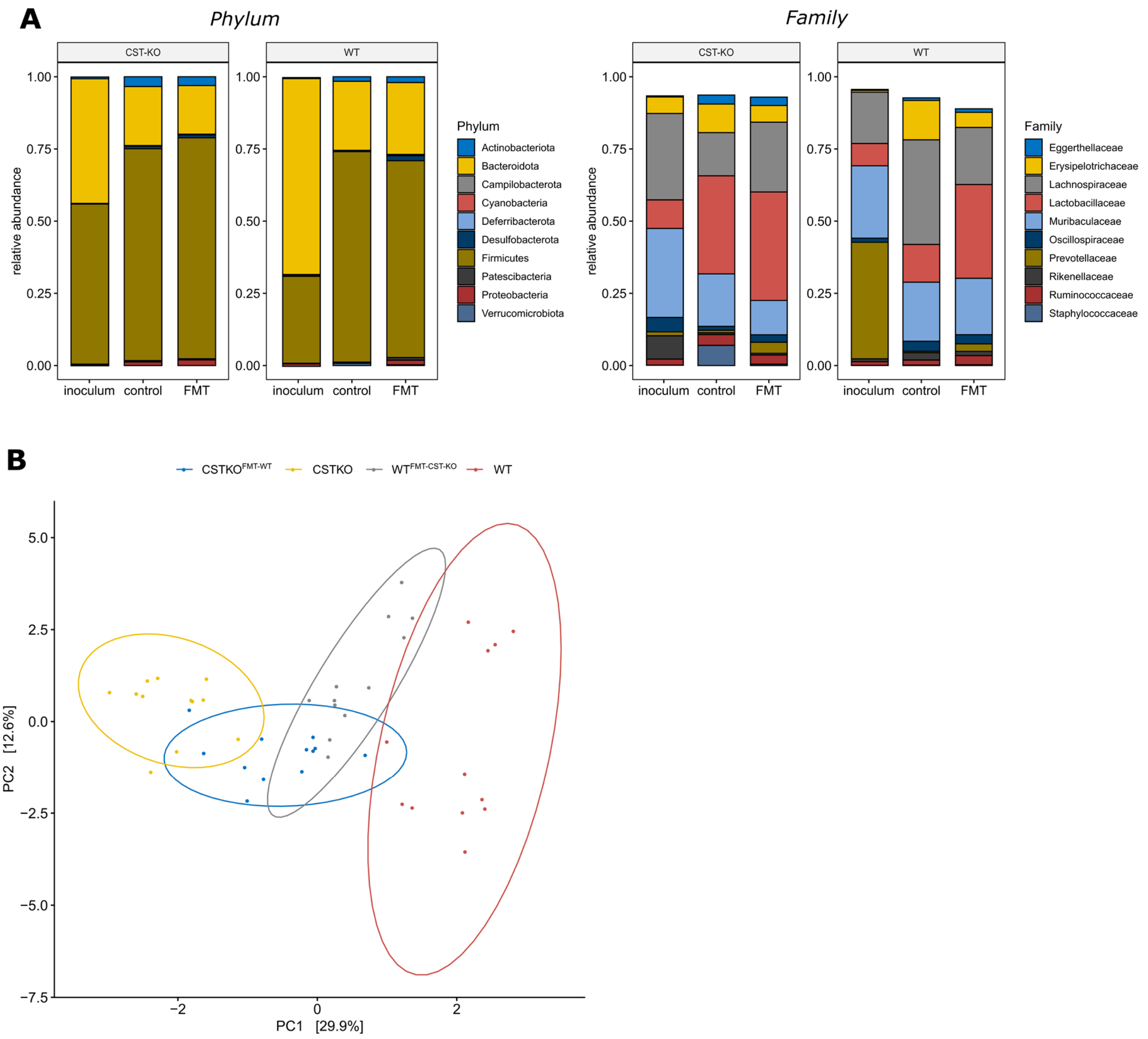


**Supplementary Figure 1.** Principal component analysis (PCA) of genus-level collapsed abundance data, shows distinct clustering of the different groups. Especially the control groups are separated, while the FMT treated groups converge to a middle point. Ellipses represent normal data ellipses produced by methods from the R package ggplot2.

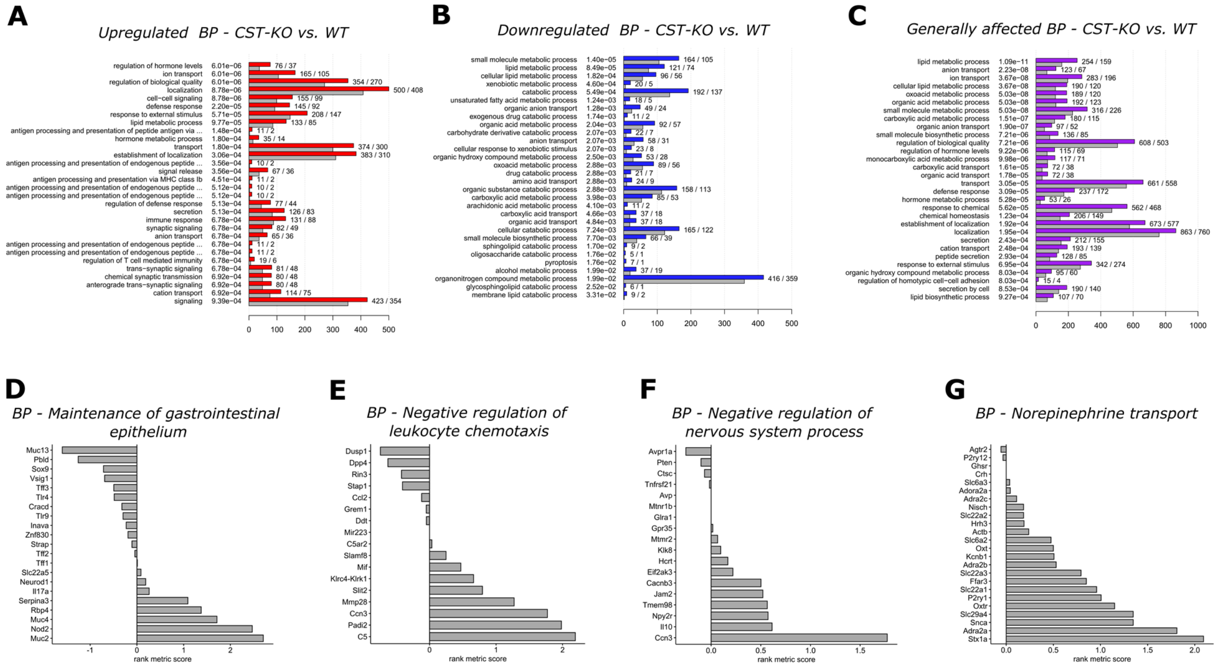


**Supplementary Figure 2**. Bar plots of the 30 most significant GO bioprocesses identified by ClusterProfiler for the comparison between WT and CST-KO control groups**.** As input the list of DEGs ($p_{adjusted}$ < 0.01) was either subset to only contain upregulated **(A)** or downregulated **(B)** genes or not filtered at all **(C)**. Coloured bars indicate the number of found DEGs in a specific category, while grey bars indicate the amount of DEGs to be expected by chance in each category. The q-values are indicated after each process (in the middle) and numbers at the end of the bars indicate the number of observed and expected genes. **(D-G)** Selection of bioprocesses, which were identified by GSEA to be significantly enriched in either genotype. Values indicate the ranking metric (signal2noise ratio) for each gene contained in the bioprocess. Positive values designate enrichment in CST-KO and negative values in WT mice respectively.

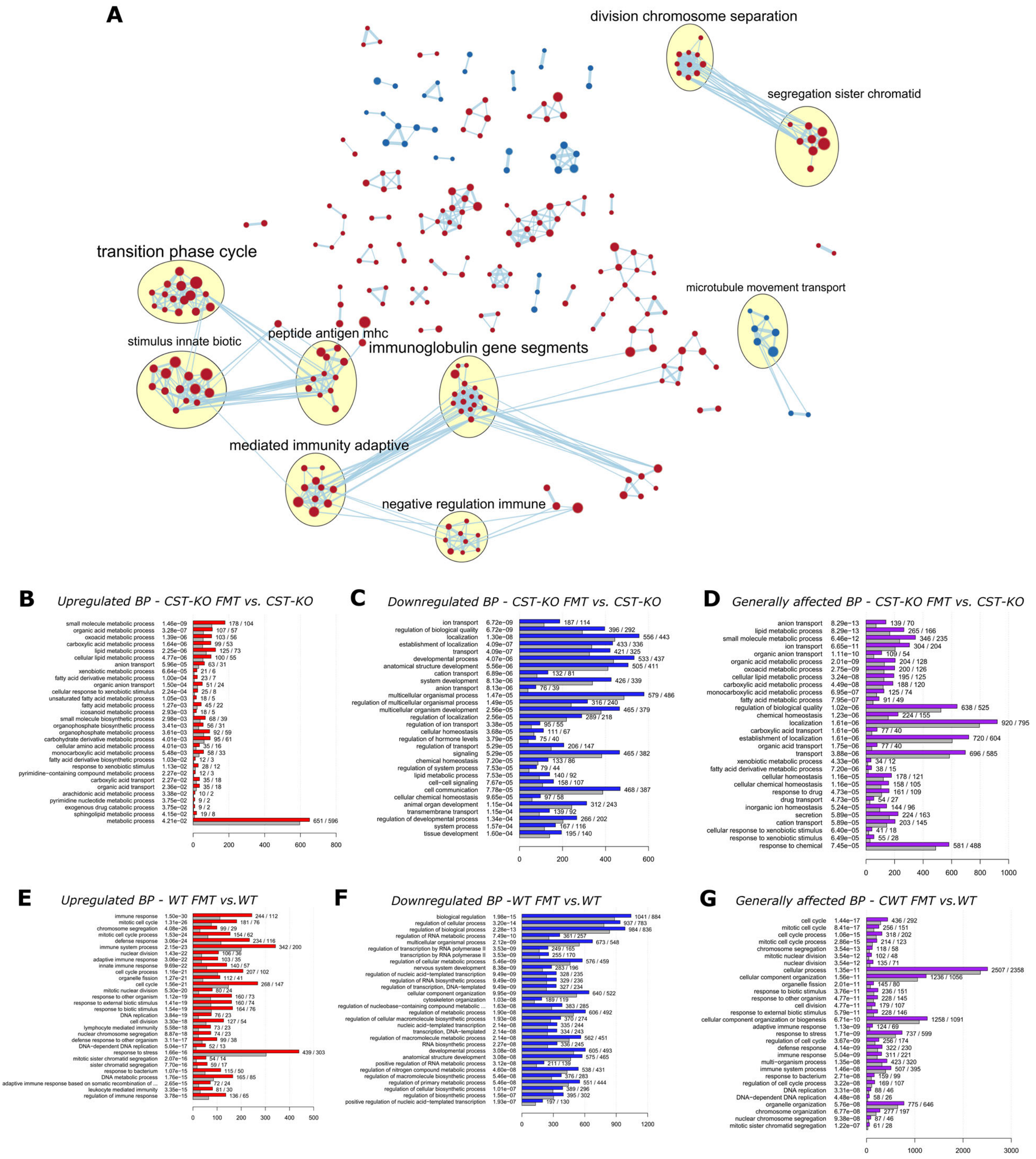


**Supplementary Figure 3. (A)** Enrichment map of significantly expressed gene sets (GO terms; nominal p-value <0.05) identified by GSEA between WT and WT$^{FMT\text{-}CST\text{-}KO}$. Blue indicates enrichment in WT, while red indicates enrichment in WT$^{FMT\text{-}CST\text{-}KO}$. Clusters were annotated with Cytoscape AutoAnnotate plugin. Connections between nodes indicate more than 25% of shared DEGs, while edge thickness correlates with the shared fraction. **(B-G)** Bar plots of the 30 most significant GO bioprocesses identified by ClusterProfiler for the comparison between CST-KO and CST-KO FMT-WT and between WT and WT FMT-CST-KO. As input the list of DEGs ($p_{adjusted} < 0.01$) was either subset to only contain upregulated **(B, E)** or downregulated **(C, F)** genes or not filtered at all **(D, G)**. Coloured bars indicate the number of found DEGs in a specific category, while grey bars indicate the amount of DEGs to be expected by chance in each category. The q-values are indicated after each process (in the middle) and numbers at the end of the bars indicate the number of observed and expected genes.

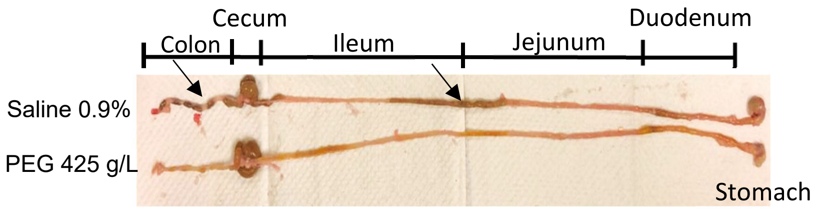


**Supplementary Figure 4** . Representative photograph showing the gastrointestinal tract of a mouse after four bowel cleansings using 0.9% saline (up) and polyethylene glycol PEG (down). Arrows depict the luminal content in the bowel still present after using saline solution but not in the PEG treated group.

**Supplementary Tables**

**Supplementary Table 1:** Clustering results from core network analysis

| Custer | No. of Nodes | Node Density | Cluster Quality | p-value | Included Nodes | Functional enrichment |
|---|---|---|---|---|---|---|
| 1 | 13 | 0.628 | 0.742 | 1.84E-05 | *Nkg7, Cd7, Cd8b1, Gzma, Ctsw, Cd8a, Cd3g, Itk, Ccl5, Hck, Lck, Il2rb* | Adaptive Immune Response, Leucocyte activation, T-cell differentiation |
| 2 | 9 | 0.778 | 0.778 | 2.11E-04 | *Uqcr11, Etfb, Atp5k, Ndufs6, Cox7b, Atp5g1, Uqcrq, Ndufa4, Dbi* | Generation of precursor metabolites and energy, Electron transport chain, ATP metabolic process |
| 3 | 7 | 0.619 | 0.722 | 0.0016 | *Mttp, Abcg8, Cyp27a1, Abcg5, Mylip, Slc51b* | Cholesterol homeostasis, Organic hydroxy compound transport, Cholesterol efflux |
